# Developmental transcriptomics of the brittle star *Amphiura filiformis* reveals gene regulatory network rewiring in echinoderm larval skeleton evolution

**DOI:** 10.1101/166991

**Authors:** David Dylus, Liisa M. Blowes, Anna Czarkwiani, Maurice R. Elphick, Paola Oliveri

## Abstract

Amongst the echinoderms the class Ophiuroidea is of particular interest for its phylogenetic position, ecological importance, developmental and regenerative biology. However, compared to other echinoderms, notably echinoids (sea urchins), relatively little is known about developmental changes in gene expression in ophiuroids. To address this issue we have generated and assembled a large RNAseq data set of four key stages of development in the brittle star *Amphiura filiformis* and a *de novo* reference transcriptome of comparable quality to that of a model echinoderm - the sea urchin *Strongyloncentrotus purpuratus*. Furthermore, we provide access to the new data via a web interface: http://www.echinonet.eu/shiny/Amphiura_filiformis/. With a focus on skeleton development, we have identified highly conserved genes associated with the development of a biomineralized skeleton. We also identify important class-specific characters, including the independent duplication of the *msp130* class of genes in different echinoderm classes and the unique occurrence of spicule matrix (sm) genes in echinoids. Using a new quantification pipeline for our *de novo* transcriptome, validated with other methodologies, we find major differences between brittle stars and sea urchins in the temporal expression of many transcription factor genes. This divergence in developmental regulatory states is more evident in early stages of development when cell specification begins, than when cells initiate differentiation. Our findings indicate that there has been a high degree of gene regulatory network rewiring in the evolution of echinoderm larval development.

**Data Depositions:** All sequence reads are available at Genbank SRR4436669 - SRR4436674. Any sequence alignments used are available by the corresponding author upon request.

## INTRODUCTION

A fundamental question in evolutionary biology is how complex characters originate. Complex structures, such as the endoskeleton, sensory organs or central nervous system, are built during animal development and encoded by a precise program(s) that requires coordinated expression of many genes regulated by large networks. A comprehensive theory formulated a decade ago by Davidson and Erwin (2006) explains both macro and micro evolutionary transitions as changes in gene regulatory networks (GRN) or rewiring (Davidson & Erwin 2006). Therefore, comparative studies of gene expression during development have been used fruitfully in locating GRN rewiring that occurred during evolution (Israel et al. 2016).

The calcite endoskeleton of echinoderms provides an ideal system to study the evolution of complex characters at the level of GRNs. The phylun Echinodermata comprises five extant classes with well-supported phylogenetic relationships, which group echinoids (sea urchin) and holothuroids (sea cucumber) (Echinozoa) as a sistergroup to asteroids (sea star) and ophiuroids (brittlestar) (Asterozoa), with crinoids (sea lilies) an outgroup (Cannon et al. 2014; Telford et al. 2014; O’Hara et al. 2014). While all echinoderms have calcitic skeleton as adults, only ophiuroids and echinoids develop an elaborate skeleton as larvae. In contrast, the larvae of the other three classes either develop only miniature ossicles, called spicules (holothuroids), or do not form a skeleton at all (McCauley et al. 2012; McIntyre et al. 2014). This provides an ideal evolutionary context to study appearance and/or reduction/loss of complex morphological characters. The most comprehensive GRN model so far studied for an animal describes the development of the larval skeleton in the sea urchin *Strongyloncentrotus purpuratus* (Rafiq et al. 2012, 2014; Oliveri et al. 2008). It explains how in the course of development dozens of regulatory genes act together to specify a mesodermal cell population, which later form two ventro-lateral clusters on each side of the primitive gut (archenteron) and finally secrete the calcitic endoskeleton typical of the sea urchin pluteus larva (reviewed in McIntyre et al. 2014). Interestingly, whereas around 30 transcription factors (TF) and a few signalling pathways are sufficient for the initiation, progression and maintenance of this process (Oliveri 2008), more than 800 genes participate in the final step of cell differentiation and biomineralization of organic matrix. These differentiation genes have been identified using transcriptomic and proteomic experimental strategies (Rafiq et al. 2014; Mann et al. 2008, 2010; Barsi et al. 2014), although their role and GRN linkages are largely un-explored. The extensive level of detail of the sea urchin GRN underlying skeletogenesis provides a useful framework to address questions about the evolution of development mechanisms through comparison with other echinoderms. Indeed, expression data are already available for a few orthologs of sea urchin skeletogenic transcription factor genes that have been identified in representatives of all echinoderm classes except crinoids (Dylus et al. 2016; McCauley et al. 2012; McCauley et al. 2010; Seaver & Livingston 2015). However, there has been relatively little comparative analysis of genes involved in skeletal differentiation in echinoderms.

Recently, biological and evolutionary studies have been transformed by immense technological improvements in sequencing technology (Goodwin et al. 2016). Relevant to this study, RNA sequencing is now an established technique that provides a practical and cheap alternative to whole genome sequencing (Wang et al. 2009) because it allows rapid advancements in molecular genetic analysis of organisms for which limited or no genomic data are available but which are of great interest from an evolutionary and/or developmental perspective. Importantly, RNA sequencing enables a global quantitative analysis of gene expression at specific stages of life and/or in particular tissues/organs. In this way it is possible to reconstruct the timeline of expression of each individual gene and determine the progression of regulatory states, which is a key first step when analysing gene regulatory networks (Materna & Oliveri 2008).

The large amount of molecular genetic information in echinoids compared to other echinoderm classes can be attributed to the fact that sea urchins have been studied extensively for over 100 years. Furthermore, the genome of the sea urchin *Strongylocentrotus purpuratus* was sequenced 11 years ago (Sea Urchin Genome Consortium 2006) and together with several improvements and additional mRNA sequencing data provides a very high quality resource (Tu et al. 2012, 2014). So far within the echinoderms, only the *S. purpuratus* genomic resources are of a high standard, although many additional species have been sequenced at lower quality (Cameron et al. 2009). Very recently the genome sequence of the Indo-Pacific sea star *Acanthaster planci* was published (Hall et al. 2017). Furthermore, transcriptomic data are available for several echinoderm species, but with significant variation in sequencing depth and quality and with most datasets limited to a single life-stage or tissue (Janies et al. 2016; Israel et al. 2016; Vaughn et al. 2012).

Within the echinoderms, the brittle star class has received growing attention in recent years (Purushothaman et al. 2014; Burns et al. 2011; Czarkwiani et al. 2013; Dupont et al. 2010) due to their phylogenetic position as a sister group of sea stars, their mode of development and regenerative capabilities. For instance, brittle stars develop a skeleton in the larvae similar to sea urchin (Dylus et al 2016; Primus 2005) and are thus a valuable model for addressing questions relating to differences and conservation of developmental genes involved in the formation of the larval skeleton. With this perspective, a single stage transcriptome identified many orthologs of sea urchin skeletogenic genes in a brittle star species (Vaughn et al. 2012), but no quantitative data on the dynamics of gene expression was provided. Furthermore, a comparison of skeletogenic regulatory states between an echinoid and an ophiuroid identified differences and similarities in the specification of the skeletogenic cell lineage (Dylus et al. 2016). Additionally, brittle stars regenerate their arms as part of their self-defence mechanism (Dupont and Thorndike 2006). The re-development of the skeleton has been characterized in detail with respect to morphology and gene expression during various phases of regeneration (Biressi et al. 2010; Czarkwiani et al. 2013, 2016; Burns et al. 2011; Purushothaman et al. 2014). Finally, brittle stars are used as important indicator species for ocean acidification studies (Dupont et al. 2010).

Here we present a *de novo* transcriptome for the brittle star *A. filiformis* obtained using four key stages of development, with the aim to provide a global quantitative assessment of developmental gene expression. We devised a computational strategy to generate a high-quality reference transcriptome, supported by several quality measures, and a reliable quantitative gene expression profile, validated on several candidates with other gene expression profile platforms, such as quantitative PCR and Nanostring. Focusing on the distinct feature of larval skeleton evolution within echinoderms, we assess the conservation of gene content by a large-scale comparison of our transcriptome with sequencing data from an asteroid, a crinoid and an echinoid. Our results reveal a high-degree of conservation of genes associated with skeleton formation in the four species, consistent with the fact that all classes of echinoderms have a well-defined adult skeleton that originated at the base of the *phylum*. Contrary to previous studies, we identify major differences in the temporal expression of regulatory genes, which suggests a high-degree of re-wiring for the developmental GRN. Furthermore, applying a fuzzy clustering approach, we find that most skeletogenic differentiation genes exhibit an increasing trajectory of expression during development consistent with their hierarchical position as the final tier of a GRN. We also present an R-shiny application to allow access to all of the data presented here for future analysis.

## RESULT

### Assembly of a reference transcriptome for *A. filiformis*

Given the similarity of development between sea urchin and brittle star (Dylus et al. 2016; Primus 2005) we performed a global comparative analysis of the gene complement and gene expression profiles of representatives of these two classes of echinoderms. To enable this, we characterize for the first time the expression of genes in the brittle star *A. filiformis* using RNA-seq technology at four chosen key developmental stages that extend over the entire development of the larval skeleton, from early cell specification to final cell differentiation. The developmental stages are: end of cleavage stage (9 hours post fertilization; hpf), a hatched blastula stage (18 hpf), 3 samples for mesenchyme blastula stage (27 hpf) and a late gastrula stage (39 hpf) (Fig. 1A). For the sequencing, we multiplexed the 6 samples using 100bp paired-end reads on 2 lanes of Illumina HiSeq 2500, resulting in ~100Million reads per sample (STab. 1 and SFig. 1). Joining all the reads from each sequenced sample and following the khmer-protocols v0.84 (Brown et al. 2013)	, we assembled a reference transcriptome that would reflect all protein coding genes expressed in the analysed stages (Fig. 1B). In this three step assembly, we first trimmed all reads for Illumina adapters and low quality basepairs, then applied digital normalization to remove overrepresented reads and erroneous k-mers (Brown et al. 2012) and finally used the resulting reads as input for Trinity (Grabherr et al. 2011) (STab. 1). Our initial assembly resulted in 629,470 sequences (N50: 1094). To determine whether the digital normalization step introduced artefacts, we assembled each individual sample omitting this step and compared with the combined assembly. We recovered over 94% of sequences using a BLASTn search (e-value 1E-20) of each individual assembly against the combined assembly (SFig. 2). Thus, we concluded that the digital normalization step did not introduce any significant bias in the combined assembly.

**Figure 1.**
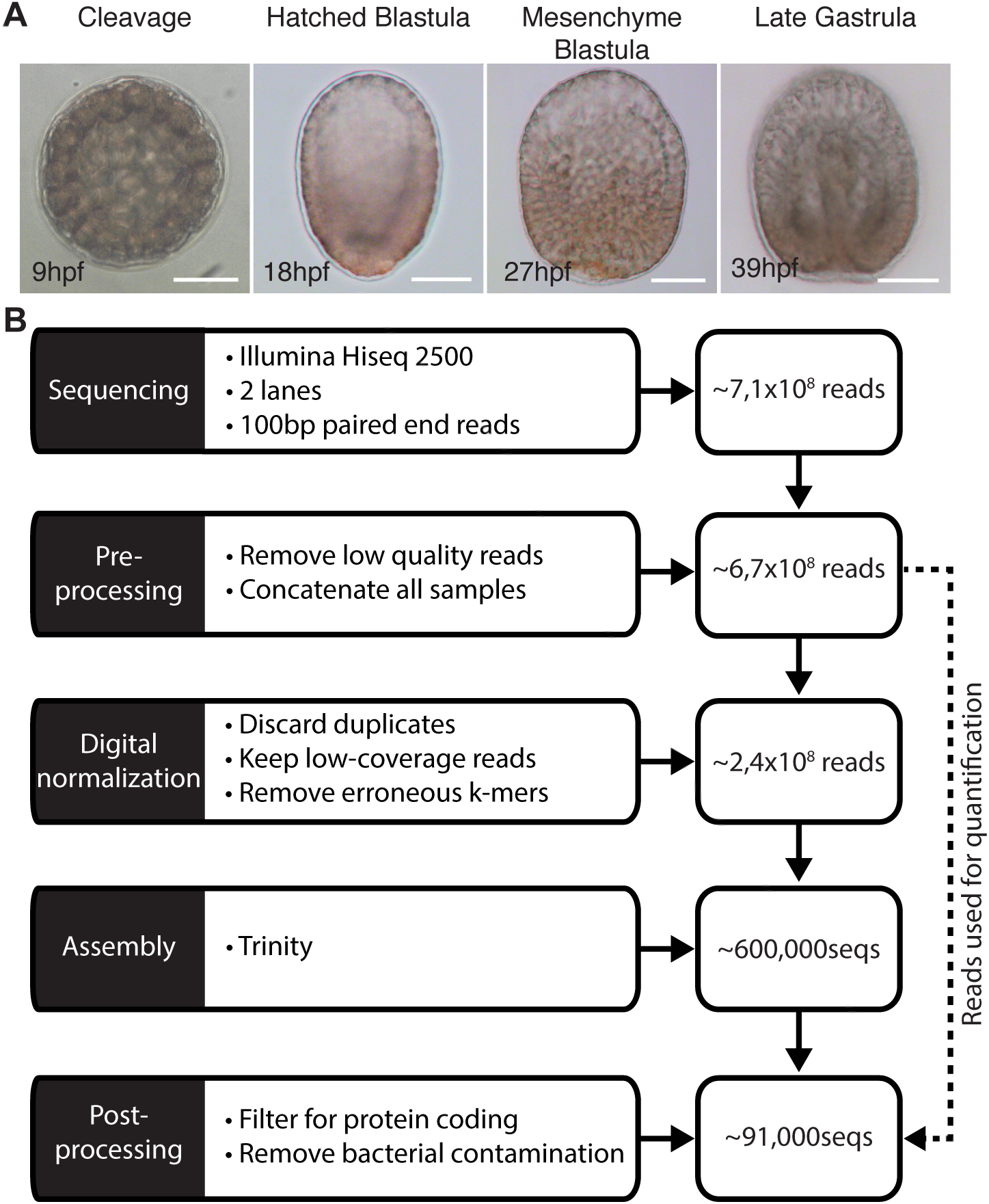
Pipeline used to obtain the *A. filiformis* developmental transcriptome. (A) Developmental timepoints used for RNA-seq: 9hpf corresponds to a late cleavage stage, 18hpf to a blastula stage, 27hpf to a mesenchyme blastula stage and 39hpf to a late gastrula stage. (B) Assembly pipeline showing the individual steps and the reduction in sequences.

Because the focus of this study was on protein-coding transcripts, we filtered our initial combined assembly for all open reading frames that have an uninterrupted coding region longer than 300bp (=100aa) using the TransDecoder package (Haas et al. 2013). This reduced our dataset to 92,750 protein coding sequences. We further removed any potential bacteria contaminates through application of a BLASTx search against 12,537,847 bacterial proteins (Uniprot DB; Bacteria Release 2014_06; 2563 species) and crosschecked the identified sequence for closer percentage of identity with hits obtained using a BLASTx (both e-value: 1E-20) search against the Uniprot SwissProt DB (Release 2014_07). Finally, we were left with 91,311 contigs (N50: 1410) constituting our reference transcriptome (RefTr) (Tab. 1). The number of contigs produced by *de novo* transcriptome assemblers is typically large as assemblers cannot differentiate between isoforms or alternative transcripts of the same gene and thus report each separately (reviewed in Moreton et al. 2014). Moreover, artefacts such as repeats, sequencing errors, variation in coverage or genetic variation within a diploid individual, create contigs that are not truly representative of different isoforms. As a result, transcriptome assemblers often report repeated contigs that differ only by a single nucleotide polymorphism (SNP), indel or fragmented versions of a transcript (reviewed in Moreton et al. 2014). Moreover, simulation studies using error-free reads showed that *de novo* assemblers inevitably produce multiple contigs for the same gene (Vijay et al. 2013). To account for this type of variation in the absence of a reference genome, but without losing sequences, we partitioned similar contigs that differ due to SNPs or indels into transcript families that share a protein identity of at least 97%. On average this approach grouped 1.3 contigs to each transcript family, resulting in 67,945 total transcript families. Unfortunately, splice variants and other artefacts are not incorporated into this type of clustering, leading to a number still larger than expected when comparing with the gene set of the sea urchin *S. purpuratus* gene set (~21.000; Tu et al 2012), the only echinoderm for which high quality genome sequence data was available when this study was conducted.

**Table 1.** Summary of quality statistics for the transcriptomic and genomic dataset used.

| Species | N25 | N50 | N75 | Longest | Mean | Median | Shortest | # Contigs |
| --- | --- | --- | --- | --- | --- | --- | --- | --- |
| *Strongylocentrotus purpuratus* | 6,297 | 4,108 | 2,438 | 22,850 | 2,821 | 2,139 | 400 | 29,072 |
| *Amphiura filiformis* | 2,601 | 1,410 | 639 | 18,993 | 925 | 525 | 297 | 91,311 |
| *Patiria miniata* | 1,131 | 474 | 351 | 23,898 | 524 | 369 | 63 | 263,867 |
| *Antedon meditarrenea* | 1,508 | 443 | 186 | 36,836 | 302 | 156 | 100 | 607,455 |

To test the quality of our assembly, we compared our RefTr with 48 isolated clones containing coding (cumulative length of 32,769bp) and UTR regions (cumulative length of 7,091bp) sequenced using Sanger sequencing technology. Using BLASTn and collecting only the top hits, we obtained an average percentage of identity of 98.6%. On an average alignment length of 588bp we found ~7 mismatches in coding sequence, resulting in an average polymorphism in coding sequences of 1.2%, a value to be expected based on the fact that clones were obtained from various batches of cDNA that are different from the samples used for the RefTr. In conclusion, we produced a high-quality reference transcriptome assembly that will provide a valuable resource for future studies in brittle star biology.

### Gene content of *A. filiformis* based on analysis of the developmental transcriptome

In order to have meaningful comparative analysis of gene expression between brittle star and sea urchin clades, which diverged roughly 480 mya (O’Hara 2014), we first classified and annotated the gene content of our RefTr and then assessed the evolutionary conservation of genes in the Echinodermata to better understand at a global level the conservation of genes and appearance of novel genes.

For this aim, and to be as comprehensive as possible, we applied independent search methods. First, we used the Blast2GO tool (Conesa & Götz 2008) that assigns gene ontology terms to each contig. Blast2GO first uses a BLASTx search (evalue: 1e-3) against the GenBank non-redundant database and this search resulted in hits for 62,388 Afi contigs corresponding to 26,010 unique genes from 1,334 different species. Consistent with ophiuroids being echinoderms, most hits were found for *Strongyloncentrotus purpuratus* (25,882/62,388 contigs), followed by the hemichordate *Saccoglossus kowaleskii* (SFig. 3). The second step of the Blast2GO pipeline performs an InterProScan to find regions within contigs that have conserved protein-coding domains. This step found 66,071 contigs with at least one region that has a recognizable protein domain. The combination of the BLASTx and interpro searches was then used to assign gene ontology terms, which provided functional classifications for 27,923 of our contigs (SFig. 3).

To proceed with a general assesment of the evolution of gene content specifically in the Echinodermata, we collected in addition to the ophiuroid *A. filifirmis* transcriptome (this study) representative datasets from the draft genome sequence of the asteroid *Patiria miniata* (Pmi) (Baylor College of Medicine: HP081117-HP139664), the genome sequence of the euechinoid *S. purpuratus* (Spu) (Tu et al. 2012; Sea Urchin Genome Consortium 2006), and the transcriptome of the skeleton-rich adult arm of the crinoid *Antedon mediterranea* (Ame) (Elphick et al. 2015) (Fig. 2A). Differences in samples, sequencing technology and assembly strategy make gene content comparisons from different species difficult. Therefore, we computed quantity and quality metrics allowing us to make meaningful statements in relation to the properties of the individual datasets (see STab. 2 and 3; SFig. 4). Importantly, at the time of the study only the sea urchin dataset had a well-curated genome and was improved by additional deep coverage transcriptome data (Sea Urchin Genome Consortium 2006; Tu et al. 2012) and is thus used here as reference for comparative analysis. Our analysis indicated that all datasets are of comparable high quality and based on our tests can be ranked from lower to higher quality as follows: Ame, Afi, Pmi and Spu (see STab. 2 and 3; SFig. 4).

**Figure 2.**
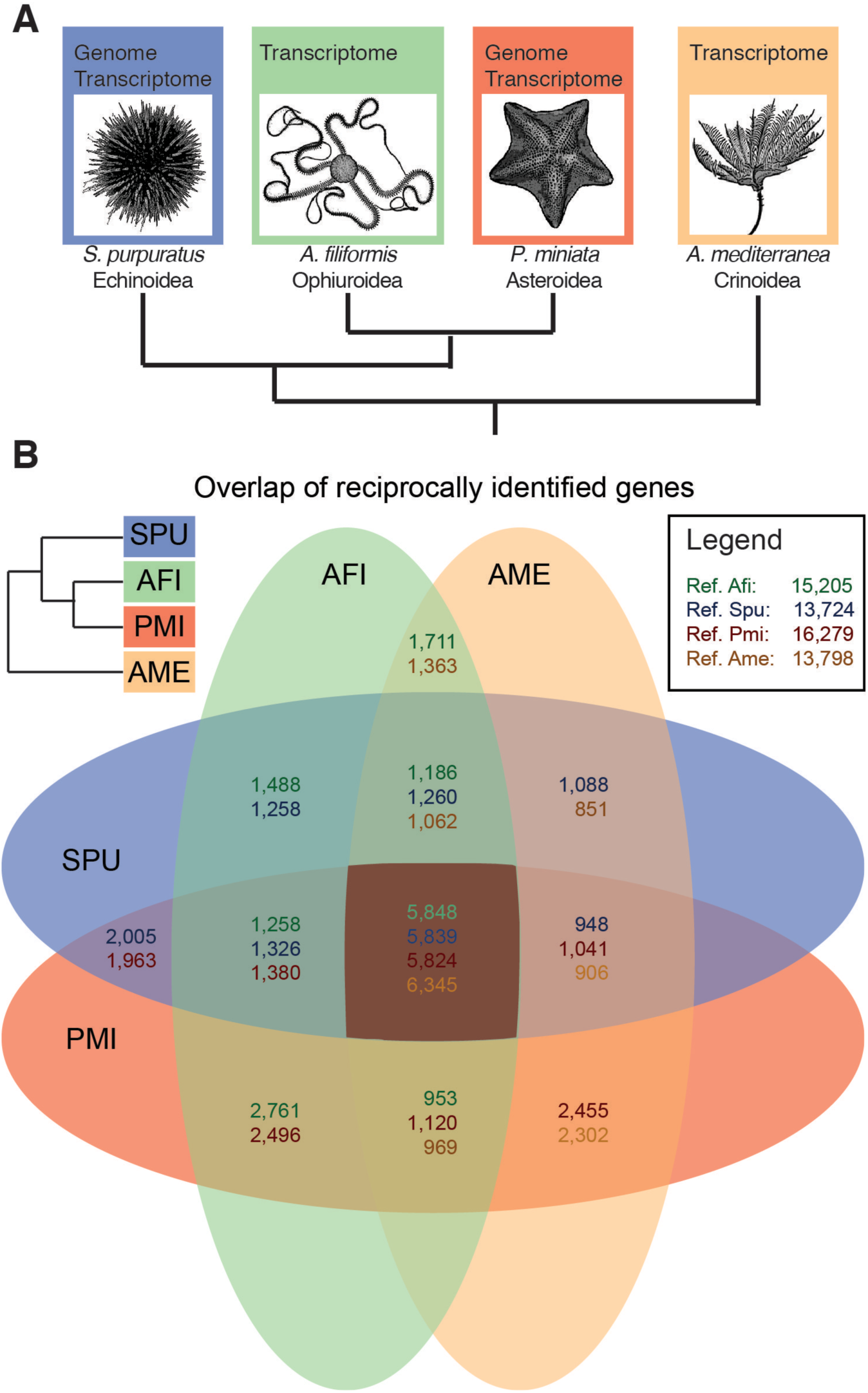
Gene content in representatives of four echinoderm classes. (A) Phylogenetic relationships of the four species compared in this study according to the currently most supported phylogeny for the classes these species belong to. (B) VENN-diagram showing the overlaps of genes that were identified using a reciprocal tBLASTx (e-value 1e-6) strategy. The different numbers in each overlap field indicate the species that was used as reference for the Blast search. Afi – *Amphiura filiformis*, Pmi – *Patiria miniata*, Ame – *Antedon mediterranea*, Spu – *Strongyloncentrotus prupuratus*, Echi – Echinoderm core (overlap of all four classes).

To gather information about the echinoderm-specific gene content we used a union of the Spu genesets predicted from genome and transcriptome databases (29,072) to identify genes in Afi and the other echinoderm species by applying a tBLASTx (evalue: 1e-6) search. For the identification we followed the khmer-protocols v0.84 (Brown et al. 2013). In this protocol a reciprocal blast is used on the sequences partitioned into transcript families. Reciprocally identified sequences are classified as orthologs and unidirectional identified sequences as homologs. Additionally, for contigs part of the same transcript family the BLAST result is propagated in order to ensure that the identification is consistent with the partition. Using this protocol, we found homologs of Spu proteins for ~58% of Afi RefTr sequences, for ~75% of Pmi genome and transcriptome derived contigs, but for only ~6% of Ame transcriptome-derived contigs. Detailed numbers are presented in Tab. 2. Importantly, the largest number of homologs of sea urchin proteins were identified in Pmi (16,211), followed by Afi (13,656) and Ame (12,982). This finding is consistent with the fact that the Pmi dataset is a combination of contigs derived from both genomic and a transcriptomic data, whereas the Afi and Ame datasets are derived solely from transcriptomes. As a positive control for our strategy, we searched the Spu dataset against itself and found 91% (Tab. 2) of hits with an evalue less than 1e-6. The residual 9% of protein coding sequences are likely to be highly similar sequences such as recently duplicated genes, different alleles, or potential wrongly annotated genes, which in general fail to give a clear unequivocal result using a BLAST alone approach.

**Table 2.** Homologs of sea urchin proteins in other echinoderms

| Species | Source | E-value | Reciprocal BLAST | Single BLAST | # Unique Spu sequences | # Query Sequences |
| --- | --- | --- | --- | --- | --- | --- |
| *A. filiformis* | Transcriptome | 1E-06 | 9,779 | 41,492 | 13,656 | 91,311 |
| *P. miniata* | Genome + Transcriptome | 1E-06 | 10,208 | 77,576 | 16,211 | 263,867 |
| *A. mediterranea* | Transcriptome | 1E-06 | 9,164 | 26,997 | 12,982 | 607,454 |
| *S. purpuratus* | Genome + Transcriptome | 1E-06 | 26,395 | 2,675 | 26,475 | 29,072 |

To determine the extent of sequence conservation in the echinoderm phylum we computed the overlap of contigs shared between species. Therefore, we searched reciprocally all versus all species (tBLASTx, evalue: 1E-6) using each time one of the four species as a reference (Fig. 2B). Our analysis shows that around 6,000 sequences are common to all species analysed, corresponding to 25% of the protein coding sequences of the sea urchin reference species. Any other combination of 2-3 species identified at least 1,000-2,000 shared genes. This suggests that in each class a specific subset of ancestral genes has been retained and consequently that others have been lost or have diverged beyond recognition with the methods employed here. Notably, we observed a higher number of genes to be shared between Afi and Pmi compared to other pairs of species (Fig. 2B). This is consistent with the recently published phylogenetic analysis of echinoderm relationships, in which sea stars and brittle stars are sister groups (Telford et al. 2014; Cannon et al. 2014). To validate this result, we applied the orthology matrix algorithm (OMA)_(Roth et al. 2008), which computes highly reliable groups of orthologous genes using the Smith-Waterman algorithm for sequence alignment. The set of orthologous genes obtained allowed us to clearly distinguish the differences in genes shared between species (Roth et al. 2008). Using OMA, we observe a much higher conservation between Pmi and Afi than in any other overlap of two species i.e. ~7000 orthologs compared to ~ 2000-4000 orthologs (SFig. 5). Moreover, the variation in the number of genes among species overlaps indicates a high level of evolutionary dynamicity in terms of gene conservation in the four classes of echinoderms analysed here. This is supported by the similar number of genes shared between two species and can be explained by the separation of the four classes early on in echinoderm evolutionary history (542 - 479 mya) followed by long periods of independent evolution (O’Hara et al. 2014; Pisani et al. 2012).

### Functional characterisation of echinoderm genes reveals conservation of a regulatory toolkit in echinoderms

A recent study explored in detail a developmental transcriptome of *S. purpuratus* in terms of gene content and established echinoderm specific ontology classifications (Tu et al. 2012). Our high quality RefTr and consistent data treatment allowed us to apply this ontology classification and to compare the abundance of specific functional classes with other echinoderms. We queried our three species for the identified genes that belong to sea urchin functional classes (SUFC; Fig. 3). From a total of 6,461 genes classified in 24 SUFCs we found 4,494 homologs in Afi, 4,407 in Ame, and 4,976 in Pmi. We classified the SUFCs in three categories of conservation using manually selected thresholds. In the first category of highly conserved SUFCs (avg(Afi, Pmi, Ame)>80% of identified Spu sequences), we find Cytoskeleton, Phosphatase, Signaling, CalciumToolkit, CellCycle, TF, DNAReplication, GermLineDeterminant, TranslationFactorTF (Fig. 3). SUFCs that are conserved at a lower level (intermediate: avg(Afi, Pmi, Ame) in between 70-80% of identified Spu sequences) are Histone, Metabolism, Nervous, GTPase, Kinase and EggActivation and whilst the lowest conservation of SUFCs (avg(Afi, Pmi, Ame) <70% of identified Spu sequences) is observed for Biomineralization, Immunity, Oogenesis, Defensome, ZNF, Apoptosis, Metalloprotease, Adhesion and GPCRRhodopsin (Fig. 3). Interestingly, the class of Biomineralization, GPCRRhodopsin, Histones and ZNF shows the highest level of variation between the three species (sd > 10%) and we find a high number of ZNFs only in brittle stars (Fig. 3).

**Figure 3.**
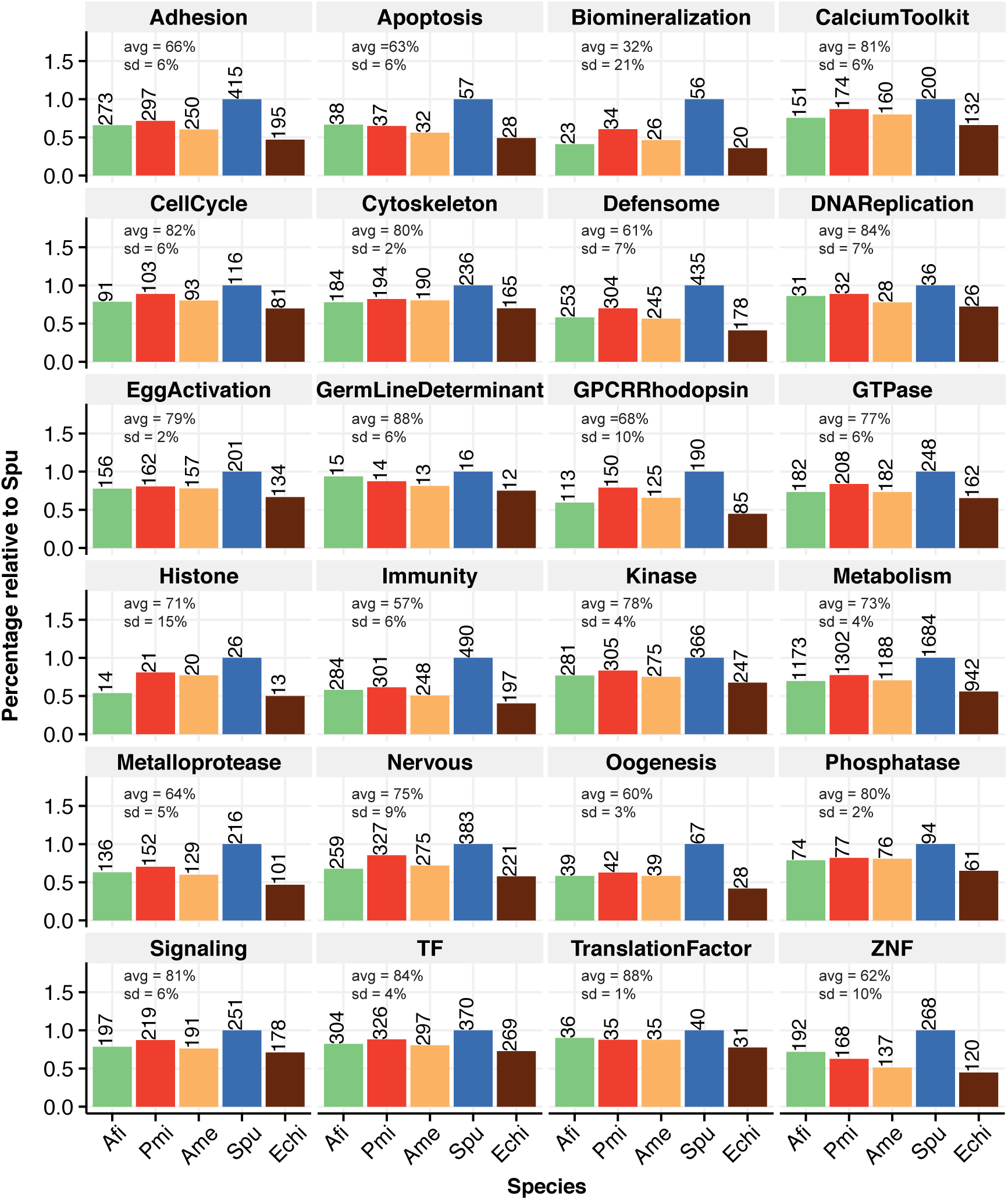
Conservation of gene functional classes in echinoderms. Sea urchin functional classes are based on *S. purpuratus* (Tu et al. 2012) and show proportions identified in the other 3 echinoderms. In the top right corner is depicted the currently most supported phylogeny for these 4 species. Average and standard deviation are calculated between Afi, Pmi and Ame and are normalized based on the sea urchin. Afi – *Amphiura filiformis*, Pmi – *Patiria miniata*, Ame – *Antedon mediterranea*, Spu – *Strongyloncentrotus prupuratus*, Echi – Echinoderm core (overlap of all four classes).

To obtain a better picture of the conservation of the developmental program in general and the evolution of the larval skeleton in particular, we focused our analysis on regulatory genes (TF and Signalling) and on biomineralization differentiation genes. Out of 368 sea urchin TF, we identified 304 in the brittle star, 297 in the crinoid and 326 in the starfish. 304 TF in the brittle star correspond to 82% of the sea urchin transcription factors (TF) and represent the cohort of TF used in this species throughout development, a number comparable to estimates obtained for sea urchin development (~80% of 283 transcription factors are expressed by late gastrula (Howard-Ashby et al. 2006)). Consistent with the fact that the starfish dataset is a combination of genome and transcriptome we find the largest number of homologs of sea urchin TFs (326) in this class of echinoderms. On the contrary, the lowest degree of conservation was observed in the crinoid (297 out of 368), which might be attributable to the fact that the Ame transcriptome was obtained from a single adult structure (the arm), although arms are formed from multiple tissue types. Generally, a similar degree of conservation was observed for signalling molecules (~76-87%), but with more variation between Pmi, Ame and Afi (Fig. 3). The high level of TF and signalling conservation indicates that echinoderms share a similar regulome.

The biomineralization SUFC shows a higher degree of variation and we find generally less genes ~41-60%, or a lower percentage of conservation. Interestingly, when looking more thoroughly in the bio-mineralization class of genes, of the 14 spicule matrix (sm) genes only one gene in Afi and one gene in Pmi were identified, indicating that the sm class of genes is significantly reduced in the Asteroidea and Ophiuroides, by comparison with the Echinoidea. Homologs of more than 50% of Spu genes belonging to the collagen, cyclophilin and carbonic anhydrase categories (STab. 4) were found all species. Interestingly, in a first assessment we found few homologs of the nine Spu msp130 genes in the species analysed here (two sequences in Afi, three in Pmi, and four in Ame), although many contigs showed sequence matches. Therefore, we investigated if there are actually more msp130 genes in the other species that Blast algorithm alone is not able to discriminate. Using 18 candidate genes, we generated a multiple sequence alignment and built a hidden Markov model (http://hmmer.org, version 3.1b) in order to scan for other contigs with a msp130 signature. With this approach, we found several candidates in our dataset that had this signature but were different in terms of their amino acid sequence. In order to investigate their relation to the sea urchin msp130 genes we built phylogenetic trees using Bayesian and maximum likelihood methods using also genes found in outgroup species. Our trees support class specific duplications of msp130 genes, as displayed by their independent expansions in different branches of the tree (SFig. 6). This analysis suggests that while all echinoderms share a similar regulome, defined as the cohort of all TF and signalling genes encoded in a genome, some classes of sea urchin biomineralization genes are either absent or duplicated independently when compared to the other 3 species analysed here.

### Skeletogenic genes are conserved within the echinoderms

All echinoderms develop a calcite skeleton and hundreds of genes are involved in this process. However, the SUFCs of sea urchin contain only 56 genes that are classified as biomineralization genes. To obtain a more precise picture of genes involved in skeletogenesis and their evolution we gathered 1006 sea urchin skeletogenic candidates based on literature searches. This extended candidate list was compiled from proteomic studies based on skeletal elements obtained from adults and larvae (Mann et al. 2010), a differential analysis of sea urchin mesenchyme blastula where skeletogenic mesenchymal cells were removed (Rafiq et al. 2014) or isolated (Barsi et al. 2014) and a large scale morpholino analysis (Oliveri et al. 2008); it is therefore representative of the skeleton developmental process from cell specification up to the deposition of the biomineralized skeleton. We updated this list with the latest annotation of the sea urchin genome and obtained 901 genes (SFile 1). Of these 901 candidates, 37 are TF and 32 are signalling molecules belonging to 5 different pathways (*i.e*. Fgf, Vegf, Delta/Notch, Wnt and BMP), whilst the rest of the genes belong to various classes of C-type lectin-type domain, carbonic anhydrases, matrix metalloproteases, known skeletogenic matrix genes (sm and msp130) and others. To maintain a very broad view, we searched the homologs of our annotated species for these candidates with the aim to find a core set of skeletogenic genes and possibly a set specifically used in the development of the larval skeleton in echinoids and ophiuroids. We find 601 candidate skeletogenic genes in *Ame*, 622 in *Afi* and 672 in *Pmi* out of 901Spu genes, which follow a trend similar to the whole gene set. To display the differences in skeletogenic gene conservation we computed the overlaps between the four species (Fig. 4). Due to the fact that skeletogenesis in the adult is a feature present in the common ancestor of extant echinoderms we wanted to check whether the 494 skeletogenic genes found in all four species are more highly conserved than a set of randomly selected genes. Therefore, we computed the overlap of 901 genes selected randomly 1,000 times and compared it with the skeletogenic gene set (SFig. 8). Our analysis suggested that genes associated with the skeletogenic process are more conserved than a set of random genes (compare 494/757 to 278/613, chi-squared proportion test p<0.001; Fig. 4 and SFig. 7). This is in line with the evolution of the biomineralized ossicle in the form of stereoms at the base of the echinoderms and a high level of conservation of this structure throughout evolution. Although, this analysis gives us a good indication of the presence or absence of genes in the different classes of echinoderms, it does not provide evidence that these genes participate in skeleton formation. Recently, using a candidate approach we showed in a multi gene expression study that of 13 TFs involved in Spu skeletogenesis 10 are active in Afi development, whilst the other 3 although expressed during development are not localized in cells giving rise to skeleton (Dylus et al. 2016). This highlights the importance to complementing transcriptomic data with spatial/temporal analysis of gene expression. Therefore, we selected from our list of 622 skeletogenic homologs 11 candidates of the differentiation cascade to investigate if they are expressed in the skeletogenic mesoderm (SM) lineage in brittle star (Fig. 4). We found that all of these genes are either expressed specifically or are enriched in skeleton-associated cells during the development of *A. filiformis*. Most of them seem to be specifically enriched in the SM lineage at late gastrula stages in cells where the skeleton is deposited. Together with our previous analysis of developmental regulatory states (Dylus et al 2016) a total of 24 genes show expression in cells associated with biomineralised skeleton conserved in two distant clades: sea urchin and brittle star. This indicates a largely similar molecular make up of calcitic endonskeleton (65%) in sea urchin and brittle star; and it is consistent with the ancient origin of the biomineralized skeleton in the form of stereom, which originated at the base of the Echinodermata phylum.

**Figure 4.**
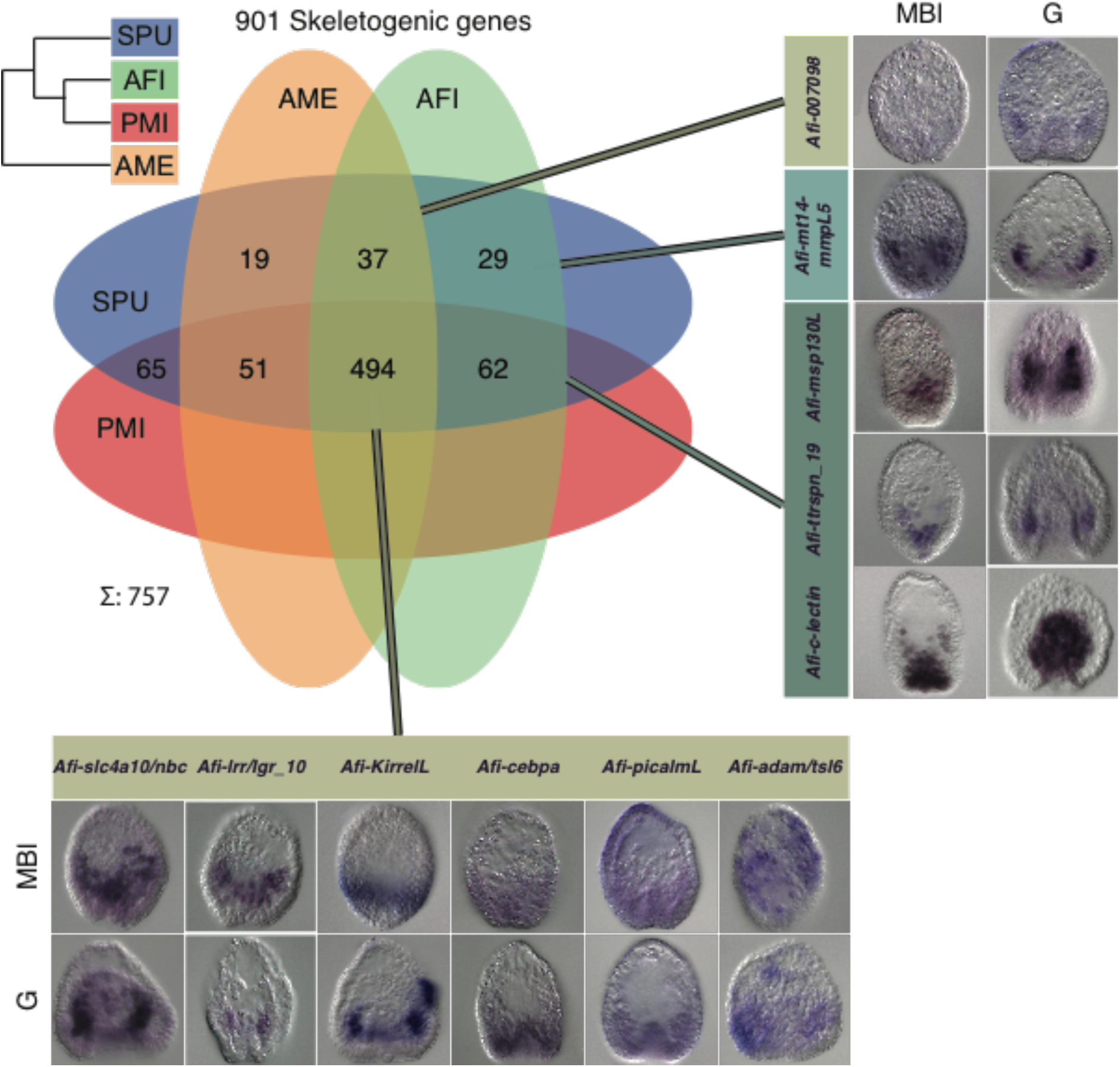
Homologs of sea urchin skeletogenic genes identified in other echinoderms and expression patterns for selected candidates. Venn diagram showing the overlap of genes involved in sea urchin skeletogenesis with homologs found in other echinoderms. 494/901 are shared between 4 classes of echinoderms, which is a higher proportion than a set of random genes (SFig. 7). Whole mount in situ expression patterns in two important brittle star developmental stages for several selected candidates from different regions of overlap reveals an association with cells associated with skeleton formation. In the top right corner is depicted the currently most supported phylogeny for these 4 species. Afi – *Amphiura filiformis*, Pmi – *Patiria miniata*, Ame – *Antedon mediterranea*, Spu – *Strongyloncentrotus prupuratus*, Echi – Echinoderm core (overlap of all four classes). MBI - mesenchyme blastula, G – gastrula.

### A quantitative developmental transcriptome for *A. filiformis* to assess the dynamics of gene expression

Our prior analysis indicates that skeleton forming genes are well conserved within the echinoderms, but what about the regulatory program? The developmental regulatory program is executed by a large GRN that tunes the expression of thousand of genes. To have an initial global assessment of the *A. filiformis* regulatory program we took advantage of the separate sequencing of four key developmental stages and the ability to obtain quantitative data from RNA-seq. While being relatively trivial to align reads when well curated gene models exists, this task is complicated for *de novo* assembled transcriptomes due to the high level of contig redundancy. In order to address this issue, we used the CORSET algorithm (Davidson & Oshlack 2014). CORSET removes sequences with less than 10 reads, which correspond to techincal background level and groups contigs to expression clusters (EC) that share the same reads, thus resulting in expression values that are equivalent to potential gene counts. In a first step this algorithm removed 9,854 sequences that were expressed with less than 10 reads. The resulting 81,457 contigs were then clustered to 37,999 ECs (min: 1seq, max: 66seq, mean: ~2.1seq per cluster; SFig. 8). In order to normalize the dataset relative to an internal standard, we computed the standard deviation for each EC between the 4 time points and selected 331 ECs with sd < 0.01 (a list of all ECs can be found in SFile 2). We then divided the RPKM corresponding to each EC by the average of the 331 ECs and multiplied each by 1 million to normalize and to obtain EC counts in transcripts per million (tpm). Because of the grouping of contigs into ECs, the previous annotation could not be directly propagated. Therefore, we associated to each EC the most frequent annotation of its containing contigs, giving orthologs priority over homologs. This caused a reduction from 13,656 to 11,695 uniquely found sea urchin sequences in Afi. Of the reciprocally identified sequences only 350 were lost during this process, resulting in 9,429 reciprocally identified sea urchin sequences. Possible reasons for this reduction are the filtering of low level of expressed sequences (less than 10 reads; see above), and contigs mapping to different genes in sea urchin actually belonging to a single one. A summary for losses mapped to the SUFCs is presented in SFig. 9. To estimate the quality of our approach we compared 29 genes quantified using qPCR and and 86 genes quantified using Nanostring in different RNA batches with the corresponding ECs. We obtained a high correlation between qPCR and ECs (r^2^ = 0.84) and between Nanostring (Geiss et al. 2008)_and ECs (r^2^ = 0.77), supporting our quantification strategy (SFig. 10 and SFig. 11). These quantitative data are now available for evaluating dynamicity of gene expression and comparative analysis and will be used for comparative gene expression with sea urchin.

### Temporal mode of TF expression in the brittle star shows many differences with the sea urchin

In order to obtain a global view of time-series expression during development and to group the genes by similar expression patterns we applied a fuzzy clustering approach (Futschik & Carlisle 2005). Based on the fact that between four time points there are 3 possible modes of expression (steady, increase or decrease) we decided to assign to each EC one of 27 fuzzy clusters (FC). This algorithm assigned 27 fuzzy clusters to the 37900 ECs. During this process 99 ECs were lost because they were not active throughout our 4 developmental time points but were expressed in one of the other two 27hpf samples that were not used for this analysis. We re-iterated this algorithm 100 times and optimized the membership of each EC to a specific FC. A closer look into the 27 FC showed four distinct modes of dynamical behavior and we decided to use this grouping for future analysis. The groups were EARLY with 10,593 FCs, INTERMEDIATE with 8,531 FCs, LATE with 9,968 FCs, and BI-MODAL with 8,808 FCs (Fig. 5 A). EARLY FCs contained ECs that showed decreasing expression across the first 3 time-points and thus likely to have a role during very early development (9hpf, end of cleavage). In these FCs we found genes that are responsible for early specification and are only transiently active. In total we found 59/287 TFs and 105/561 skeletogenic genes that showed a decreasing trajectory over the 4 time-points. In this group, only *Afi-pplx* was found as a gene involved in Afi skeleton specification. In the INTERMEDIATE group were genes whose expression trajectories peak either at 18hpf or 27hpf and then decrease steadily. Examples of genes found in this group are *Afi-alx1, Afi-tbr, Afi-gataC* and *Afi-erg*, TFs that have been shown to be expressed in mesodermal cells of the Afi embryo and known to play a role in the specification of mesoderm (Dylus et al. 2016). In total this group comprises 66/287 TFs and 68/561 skeletogenic genes. In order to form the extended larval skeleton, we expected most of the skeletogenic genes previously described to be expressed at the moment of the deposition of the calcite skeleton, and therefore to show an increasing pattern of gene expression. Indeed most of the skeletogenic genes were clustered in the LATE group 287/561. Among others, this group contained the biomineralisation genes *Afi-p19 (Cah10L), Afi-p58a, Afi-p58b, Afi-ttrspn_19, Afi-slc4a10/nbc* and *Afi-c-lectin*, all expressed in skeletogenic cells in brittle star (Fig. 3 and Dylus et al. 2016). Moreover, the LATE group contained most of the active TFs 132/287, consistent with the increasing complexity of cell types over developmental time. The final group, called BI-MODAL consists of two expression peaks throughout the 4 time-points and contains 30/287 TF and 101/561 skeletogenic genes. This group contains genes that might be expressed in different domains during development, potentially having two (or more) roles throughout development. Examples are *Afi-hesC* and *Afi-delta*, which are first expressed in the mesodermal cells at the vegetal side of the embryo at blastula stage (18 hpf) and then in scattered cells in the ectoderm at gastrula stage (39 hpf) and at the tip of the archenteron throughout gastrulation (Dylus et al. 2016). Based on the fact that our 4 time-points correspond to 4 different stages of development, our grouping shows consistent activity of TF involved in multiple stages of cell specification.

**Figure 5.**
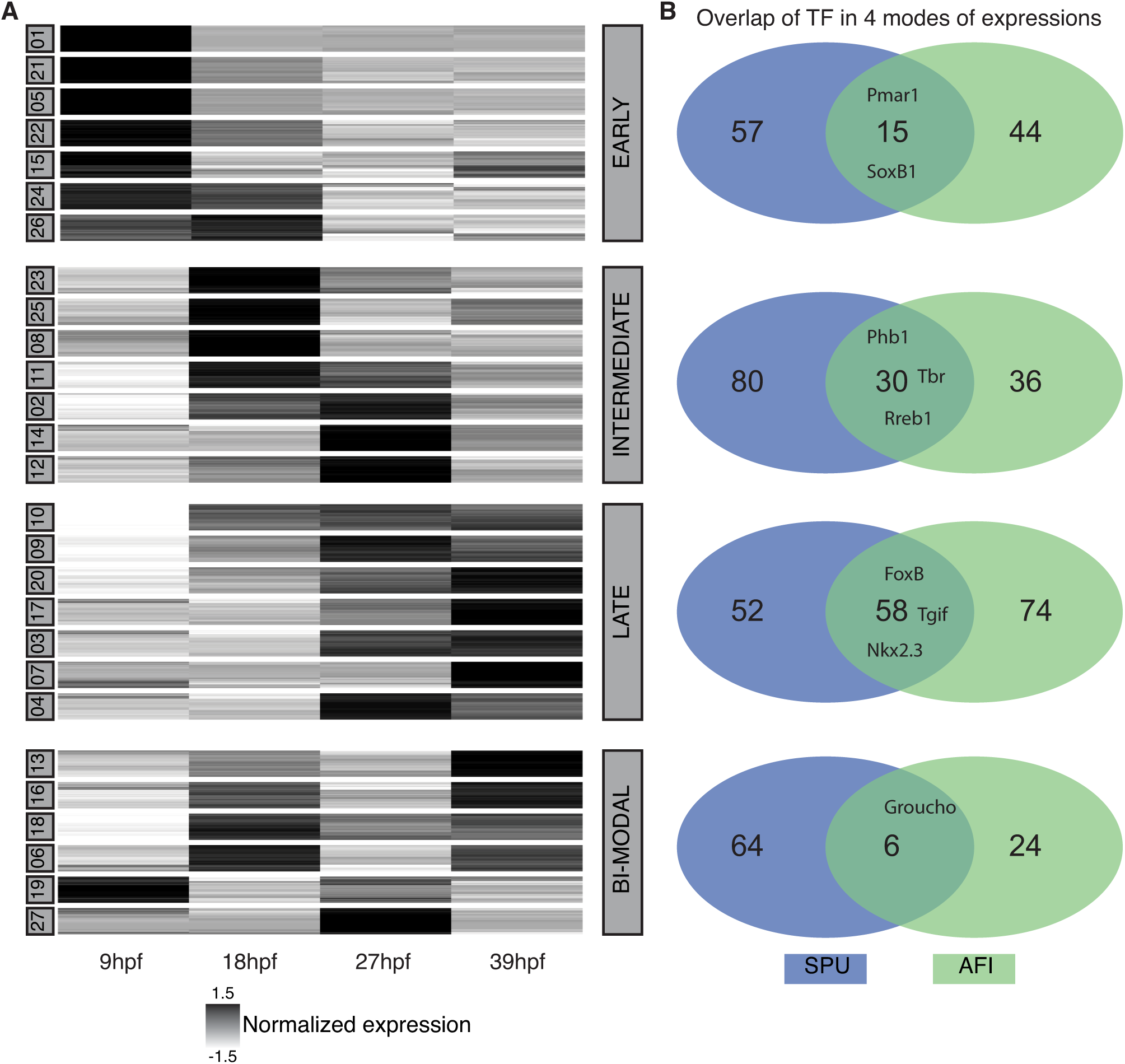
Global *A. filiformis* gene expression and comparison of larval regulatory states. (A) Fuzzy clustering of 39,000 ECs in 27 clusters of four developmental time-points sorted in four distinct modes of expression (EARLY, LATE, INTERMEDIATE, BI-MODAL). Each line represents the expression of a single gene, and the grey intensity indicates the normalized expression (B) Comparison of TF in the 4 modes of expression between sea urchin (SPU) and brittle star (AFI). The majority of TFs show differences in expression.

The direct output of a GRN is the temporal expression profile of each gene throughout time and each expression profile is linked to its regulatory state. Therefore, comparing temporal expression profiles between two species provides a first glimpse on GRN rewiring and on heterochonic gene expression. In order to evaluate the differences and commonalities of TF usage between sea urchin and brittle star, we selected 4 time points that correspond to similar stages of development from the sea urchin transcriptome: they are 10, 18, 30, 40 hours post fertilization (hpf) in agreement with the comparative developmental stages previously described (Dylus et al 2016). On these *S. purpuratus* transcriptome stages we performed a fuzzy clustering as for *A. filiformis*, and we then grouped the clusters based on the above-mentioned criteria. We identified in the EARLY category 72, the LATE 110, the INTERMEDIATE 110 and in the BI-MODAL 70. out of the 368 TFs, and six genes are not classifid due to the too low levels of expression. When comparing TF expression, and therefore the developmental regulatory states between *S. purpuratus* and *A. filiformis*, many differences emerge in the four categories. In all four categories we see more variation than overlap of TFs (Fig. 5 B). For example, only 15 transcription factors in the EARLY category are in common between the two species (e.g. *pmar1* and *soxB1*), whereas 44 Spu homologs in Afi categorized as EARLY differ from other 57 TF in Spu are expressed in this mode. Other examples of common transcription factors are for BI-MODAL *groucho*, for INTERMEDIATE *alx1, erg, foxM, mitf*, and for LATE *foxB, hnf4, tgif*. A summary of all TF can be found in SFile 3. This comparison highlights that TFs are used differently, or at least with a different timing of expression, during the development of the two species. Examples of such genes are *hesC* and *ets1/2*. Notably, there are more differences in the early phases of development when cell specification begins, than in the late stages when cells initiate the final differentiation. Given that the direct output of a GRN is reflected in the temporal gene expression, this suggests differences in the topology of gene regulatory networks between Afi and Spu.

## DISCUSSION

Here we present a *de novo* transcriptome of *A. filiformis* that samples four important stages of the embryonic development of this organism. We also present an overall strategy to effectively compare different data sets and to use RNA-seq quantitative data in the absence of a reference genome. Our data and assembly/annotation strategy is then used to obtain insights into two key evolutionary questions: how did the larval skeleton in echinoderms evolve and how conserved is the regulatory program of the pluteus larvae of sea urchins and brittle stars?

To assemble the *A.filiformis* RNA-seq data, we used a strategy with digital normalisation followed by application of the Trinity assembly. Our approach with digital normalisation allowed us to obtain a reference transcriptome that incorporated six independent samples within four weeks of computation on a server with only 64GB of memory RAM, with quality comparable to assembly obtained with non-normalised data. Our comparison is in agreement with what was observed by Lowe et al. 2014 for the assembly of sequence data from two closely related ascidians, for which a systematic comparison of assembly with and without digital normalisation showed no inclusion of computational artefacts, but a reduction of time and resources needed for the assembly. We showed that our RefTr is of high-quality by various computational and experimental methods and we also applied the computational quality control to the other datasets to strengthen the subsequent comparative analyses. In the developmental transciptome the depth of sequencing (~100M reads per sample) and the combination of samples from multiple stages were important driving factors that made such a high quality assembly possible. Altogether our analysis shows that deep sequencing combined with a good pipeline can result in an assembly that is comparable to a genome in terms of gene capture. This is illustrated by the high number of genes that showed more than 90% identity to genes in the Swissprot database. Thus, our transcriptome performed best when compared to other genome and transcriptome datasets (SFig. 4). Interestingly, our extraction of protein-coding genes reduced the total number of contigs from ~600,000 to ~90,000 (15%) increasing the N50 value, however not affecting gene recovery, as shown in the CEGMA test (STab. 3). Based on our analysis only 15% of the Ref Tr sequences are protein-coding, giving rise to a particular question: what are the residual 85% of sequences? One possibility is that they are part of non-coding sequences (e.g. non coding RNA, transcribed pseudogenes) or partially or wrongly assembled transcripts. Efforts to obtained genome sequence data for *A. filiformis* are underway to help obtain answers to these questions. Indeed, studies on human genomes show than more than 60% of the genome is reproducibly represented in long RNA sequences, while only 2.9% are represented by protein-coding sequences (The ENCODE Consortium 2012).

During the Cambrian period the rapid expansion of animal life was associated with acquisition of the capacity to form hard mineralized tissues, as testified by the first appearance of a fossil record for many phyla. Amongst others, echinoderms evolved their characteristic calcitic porous endoskeleton formed of magnesium rich calcium carbonite and occluded proteins (Bottjer et al. 2006; Gilbert & Wilt 2011). A first step towards understanding evolution and developmental genetics of a complex character such as mineralized skeleton is to perform a comparative and phylogenetic analysis of gene content (Fig. 2). For this reason we compared four echinoderm classes, 3 of the Eleutherozoa subphylum (Echinoidea, Ophiuroidea and Asteroidea) and a crinoid outgroup, with a focus on the genes involved in skeleton formation. Studies on sea urchins have shown that several genes used during adult skeleton formation are also used in larval skeleton (Gao & Davidson 2008; Mann et al. 2010), leading to the idea that an ancient regulative and differentiation module originated at the base of the phylum Echinodermata and then was secondarly co-opted to form larval skeleton. However, it is hotly debated whether this happened only once in the branch leading to the Eleutherozoa, or whether it occurred independently in both the sea urchin (Echinoidea) and brittlestar (Ophiroidea) lineages. The two transcriptomes used in this analysis correspond to stages (late gastrula, for *A. filiformis*) or structure (adult arm for *A. mediterranea*) in which the biomineralized skeleton has been deposited. Therefore, expression of genes involved in this process must be highly represented. It is important to clarify that due to nature of this comparison, genome vs transcriptome, we can unequivocally evaluate only the gene (or proteing coding transcripts) presence in at least two data set. On the other hand, in the context of skeleton development, the gene absence in *A. filiformis* and *A. mediterranea* can be interpreted as lack of expression in stages or structures with skeleton, sequenced in this study, and hence no-usage in building the skeleton of these two organisms.

Our analysis revealed a gene toolkit of 494 genes conserved in all four echinoderm classes (Fig. 4), which potentially corresponds to the echinoderm core of skeletogenic genes. Indeed our analysis of spatial expression shows that several of these genes are expressed in cells known to form the skeleton in the developing *A. filiformis* embryo (Fig. 4; Dylus et al. 2016) and for few of them also known to be expressed during *A. filiformis* adult arm regeneration (Czarkwiani et al. 2013, 2016). Of the initial 901 gene set only 37 are TFs and 32 signaling molecules. Of these regulatory genes, 84% (58/69 regulatory genes) are conserved in all the echinoderm classes analysed, while the other genes, which can generally be classified as differentiation genes, are conserved only for the 52% (436/832) in all the classes, indicating a higher conservation of the skeletogenic cells regulatory program and a rapid evolution of echinoderm skeleton-forming genes. A closer look into these 436 genes using the sea urchin functional classes revealed that metalloproteases and biomineralization genes are actually the most variable class of genes (SFig. 9). This observation indicates that solely looking into these two categories can produce a biased picture of evolution, because only these two categories of differentiation genes showed high level of variation and indicate low selective pressure. How can we explain the variation in the biomineralization genes? They are grouped in six categories of which collagens, cyclophillins, carbonic anhydrases and an unnamed category (Tu et al. 2014), which include P16 (Cheers & Ettensohn 2005) and other genes, are highly conserved in our selected representatives of the four classes of echinoderms. On the other hand, of these six categories, msp130 and spicule matrix (sm) genes show the highest level of variation. Indeed, of the nine sea urchin msp130 genes only two are found in all four species analysed (*Spu-Msp130r6* and *Spu-Msp130L*). An in depth look into the brittle star transcriptome, using a hidden Markov model, revealed also the presence of seven other msp130 contigs that show differences at the amino acid level higher than the 1.2 % of polymorphism identified in coding region, suggesting the presence of several genes. Indication of clade specific expansions took place is strongly supported by our phylogenetic analysis (SFig. 6), which shows a consistent group of sea urchin *Msp130* genes with various paralogues represented in both sea urchin species analysed (*S. purpuratus* and *L. variegatus*), a different group of Ophiuroid *Msp130s*, as well as other clade specific expansions consistent with what has already shown for *Msp130* genes in molluscs and anellids (Szabó & Ferrier 2015). Concerning the spicule matrix (sm) genes, out of the 14 genes identified in sea urchin only the C-lectin that does not contain a proline rich region is conserved in all four species. Therefore, no spicule matrix genes, characterised by a C-lectin domain and a conserved proline-rich domain (Livingston et al. 2006), are found in any other class of echinoderm in stages when skeleton is built, making them likely to be a sea urchin specific set of skeletogenic matrix genes.–Further support for this hypothesis is provided by the following observations: First, a proteomic study of skeletal elements in another species of brittle star *Ophiocoma wendtii* did not find orthologs of these genes (Seaver & Livingston 2015). However, other potential candidates of c-lectin type genes for brittle star skeletogenesis were obtained, which are also present in our transcriptome of *A. filiformis* and which are expressed during larval and adult skeletogenesis (Dylus et al. 2016; Czarkwiani et al. 2016). Second, in the *S. purpuratus* genome the sm genes are present in mini clusters of tandem repeated genes (STab. 6 and SFig. 12) suggesting a relatively recent duplication of these genes in the sea urchin lineage. Third, no such gene has been found in the hemichordate *Saccoglossus kowalevskii* (Cameron and Bishop, 2012), an outgroup of all echinoderms, confirmed by the absence of spicule matrix genes in the crinoid transcriptome analysed in this work (STab. 4). Both spicule matrix genes and msp130 genes have been highly duplicated in sea urchin, as seen in the many tandem duplications, and the presence of both in the pencil urchin *Eucidaris tribuloidis* (Cameron et al., 2009) suggests their evolution already in the common ancestor of cidaroids and euechinoids. In this context, it would be interesting in future studies analyses holothuroids as sister class to the echinoids to pinpoint more exactly the evolutionary of this category of biomineralization genes. Interestingly, similar to these findings in echinoderms, the rapid parallel evolution in different lineages of genes associated with skeleton formation has also been reported for shell genes in molluscs and brachiopods (Jackson et al. 2010; Luo et al. 2015).

The fact that both *msp130* and sm genes are expressed in both adult and larval skeletal structures in sea urchin (Mann et al. 2010) suggests that the evolution of sm genes in echinoids and the independent expansion of *msp130* genes occurred before the evolution of the echino-pluteus, the sea urchin larva with extended skeleton. Similarly, in brittle stars the specific *Afi-Msp130L* is expressed in the larval skeletogenic cells, supporting the argument that larval skeletogenesis evolved independently in the two lineages, potentially in both cases as a co-option of the adult skeletogenic program after clade-specific gene expansion took place. Other evidence in support of evolutionary divergence of the echinoid and ophiuroid pluteus larvae is provided by our comparative analysis of regulatory states in developing embryos (Fig. 5), defined as the sum of transcription factors expressed in a given cell at a given developmental time. We compared the transcription factor usage in *S. purpuratus* (Tu et al. 2012) with the one of *A. filiformis*, taking advantage of the quantitative aspects of transcriptome data and the sequence data from four key developmental stages: cleavage stage (9hpf), when maternal mRNA are still present and the zygotig genome starts to be expressed; blastula stage (18hpf), when territories that will give rise to multiple cell types are specified and transcription factor genes are express in a territorially restricted manner (Dylus et al. 2016); mesenchyme blastula (27hpf), when territories are further subdivided, cells continue in their specifaction pathway, and morphogenetic movements commence; and finally gastrula stage (39 hpf), when cell types are specified, morphogenetic movements are almost completed and cell differentiation is underway. This comparison shows that the early regulatory states, which determine the developmental GRN, of these two species are quite different. On the contrary, when cell types are specified and terminal selector genes (LATE genes in this analysis) are expressed (Hobert 2008), they show a similar regulatory make up in these two classes of echinoderms, suggesting extensive GRN rewiring in the early stages of development. Taken together, our findings are in agreement with the hypothesis that the peripheries of the GRN (i.e. early regulatory input and differentiation gene bateries) are the least constrained and thus the most frequently changed (Davidson & Erwin 2006) part of a GRN, while the phylotypic stage (identified as the gastrula stage in echinoderms; (Raff 1996; Levin et al. 2016)) is subject to strong evolutionary constraints. In this view our data support the idea that the regulatory states that define cell type identities, before differentiation, are the most evolutionary stable compared to early specification regulatory states. In the case of the developmental program for echinoderm skeleton, this likely corresponds to the transcription factors conserved in all four classes analysed here and known to be expressed in skeletal cells (Oliveri et al. 2008; Dylus et al. 2016; Czarkwiani et al. 2013). Indeed the high degree of conversation in all four classes is consistent with all echinoderms forming an adult skeleton by similar ossicle units: the stereom (Bottjer et al. 2006), and indicates that the GRN for adult skeletogenesis is a highly conserved feature. This is additionally supported by comparing expression patterns of several genes in juvenile or adult stages (Gao & Davidson 2008; Morino et al. 2012; Czarkwiani et al. 2013), which show a high degree of conservation in cells that participate in adult skeletogenesis. Additionally, in brittle star development most differentiation genes show an increasing trajectory over time consistent with their role in the final differentiation of the biomineral structure.

The modelling of developmental GRNs requires knowledge of spatial and temporal expression. For a GRN analysis comprising a few genes, the integration of such data is a relatively simple task. In a systems biology perspective, however, where hundreds or thousands of genes are considered simultaneously, it is easy to lose track of the important details of few or single genes, especially when working on novel systems with little to no access to the established data. Thus, we developed a website (http://www.echinonet.eu/shiny/Amphiura_filiformis/) using R-shiny that allows users to query different types of information, similar that implemented by Tu and collaborators in 2014 for *S. purpuratus* (Tu et al. 2014). Using the statistical programming language R as the backbone, our website provides a platform to easily query and find genes of interest. It gives access to annotations, expression levels, sequence information, differential screening and spatial expression patterns. Contigs can be queried by annotation, expression cluster id, contig id and additionally by the sea urchin functional classification. Thus, for example one can easily retrieve all transcription factors sequences and their expression temporarily and spatially (where available). Moreover, spatial expression data can be extended by simply adding a folder with the contig id and the individual pictures as JPEG files. The code is publicly available on github (https://github.com/dvdylus/Echinoderm-Web). In future work, this website will be extended with data from regenerating arms produced in our laboratory and will thus create a unique resource to establish the brittle star *Amphiura filiformis* as developmental and regenerative model system.

## METHODS

### Experimental Techniques

#### Embryological techniques

*A. filiformis* cultures were set up as previously described (Dylus et al. 2016). At the desired stage, embryos were collected for RNA extraction and/or fixed for WMISH as described in (Dylus et al. 2016).

#### Cloning and Probe synthesis

All genes used for spatial expression analysis by WMISH were PCR amplified from *A. filiformis* cDNA and cloned in pGEM-T easy vector system (Promega) or Topo PCR cloning system (Invitrogen) accordingly to the manufacturer’s instructions. Antisense probes labelled with DIG (Roche) were synthesized as previously described (Dylus et al. 2016). Primers are presented in Supplementary Table 5.

#### Quantitative PCR (qPCR)

qPCR was performed on different biological replicates to those used for the mRNA-seq, employing the procedures described previously (Dylus et al. 2016).

#### Whole mount in situ hybridization (WMISH)

Spatial expression of selected genes at mesenchyme blastula (24hpf) and (27 hpf) were characterised using WMISH as previously described (Dylus et al. 2016).

#### RNA extraction

For mRNA sequencing, embryo samples of a single male and single female culture were collected at 09hpf, 18hpf, 27hpf and 39hpf. At 27hpf three samples were collected. The RNA extraction was performed as previously described (Dylus et al. 2016). The quality of extraction and concentrations were checked using NanoDrop 2000 and Bioanalyser.

#### mRNA sequencing

Sequencing libraries were prepared using the TruSeq RNA library preparation protocol. The samples were sequenced with the Illumina v3 chemistry using the multiplex paired-end sequencing protocol. The sequencing was performed on an Illumina HiSEQ 2500 with 100bp paired-end reads. To reach optimal coverage we sequenced 2 lines multiplexing the 6 samples. Library preparation and sequencing were performed at the SickKids Hospital Toronto, Canada.

### Computational procedures

If not otherwise stated, all computational work was performed on an Apple Mac OS X 10.6 server with 24 cores and 64GB of memory.

#### Assembly

The assembly pipeline and annotation followed a set of unified protocols described in (Brown et al. 2013). The obtained reads were trimmed for adapters and for low quality sequences using Trimmomatic v0.27 (ILLUMINACLIP:Adapters.fasta:2:30:10; HEADCROP:12) (Bolger et al. 2014)	. Quality filtering was performed using the FASTX-Toolkit (v0.0.13.2; fastq_quality_filter – Q33 –q 30 –p 50). The quality filtered and trimmed reads were then digitally normalized (Brown et al. 2012). Once all filtering was completed, reads from all stages were combined and the transcriptome was assembled using the Trinity package (v2013-02-25) (Grabherr et al. 2011). Partial and complete open reading frames (ORFs) with a minimum length of 100 amino acids were predicted using the TransDecoder (version rel16JAN2014) script. Bacterial contaminants were obtained using mpiBlast (v.1.6) (Darling et al. 2003) with e-value 1E-20 and crosschecked with hits obtained against UniProtKB -SwissProt with the same e-value. Searches with mpiBlast were run on the Legion HPC cluster at UCL on at least 40 cores. Sequences with higher similarity to the bacterial database were removed from the dataset. The cleaned ORF dataset represents the reference transcriptome (RefTr). All reads were deposited in the NCBI Short Read Archive (SRA) under accession numbers: SRR4436669 - SRR4436674.

#### Preparation of other datasets

Transcriptome sequence data from *Antedon mediterranea* was obtained by the Elphick lab at Queen Mary University of London, as reported previously (Elphick et al. 2015). To obtain a complete picture of coding sequences from *Patiria miniata*, we combined both genomic derived coding sequences and transcriptome sequences from http://echinobase.org.

#### Quality Assessment

Completeness of our transcriptome was estimated using CEGMA (v2.5) (Parra et al. 2007). Full-length distributions were estimated by considering all unique hits determined by BLASTx (1e-20) against the UniProtKB-SwissProt database and application of scripts included within the Trinity application.

#### Annotation

All BLAST (Altschul et al. 1990) searches were performed using a local NCBI-BLAST (v2.2.25) with e-value of 1e-6. The RefTr was annotated against the sea urchin *S. purpuratus* transcriptome sequences and against the UniProtKB-SwissProt database. One directional BLAST identified presumed homologs and reciprocal BLAST identified presumed orthologs. Gene Ontology classification was performed based on a previous sea urchin specific classification (Tu et al. 2012). For consistency purposes sequences obtained for the starfish *P. miniata* (echinobase.org) and the crinoid *A. mediterranea* raw sequences (Elphick et al. 2015) were annotated using the same combination of one-directional and reciprocal BLAST (e-value 1e-6) against the sea urchin transcriptome database.

#### Abundance estimation

The quality filtered trimmed reads were re-aligned on the reference transcriptome using bowtie (v0.12.9) (Langmead 2010) with parameters set as in RSEM (Li & Dewey 2011). The bowtie output was loaded into CORSET in order to obtain counts for clusters of contigs that shared reads, rather than individual contigs (Davidson & Oshlack 2014). This is equivalent to a potential “gene” count adding up all “isoform” counts. Normalization by internal standard was performed as follows: First, individual clusters were normalized by their peak of expression in the time-course data (9hpf, 18hpf, 27hpf and 39hpf); then, for each cluster the standard deviation was calculated and clusters with standard deviation below 0.01 were chosen as internal standard; and finally, an average of these clusters was used as normalization factor and each cluster was divided by this normalization factor and multiplied with 1,000,000. All downstream analysis was performed using customized R and bash scripts. In order to make statements about annotation content in the individual clusters, the most frequent annotations for each expression cluster were considered.

#### Expression clustering of time-series data

To sort expression clusters by their individual trajectories we applied the fuzzy clustering algorithm (Futschik & Carlisle 2005). We used 27 fuzzy clusters, based on the assumption that between 4 sampled time points the expression either increased, decreased or did not change giving 3^3^ (27) possibile paths for each trajectory. Note here the difference between a fuzzy cluster and an expression cluster: a fuzzy clusters describes a group of expression clusters that share similar trajectories over time. Since fuzzy clustering does not allocate each transcript always to the same cluster, we re-iterated this algorithm 100 times to find for each expression cluster the most probable fuzzy cluster membership.

#### Estimation of phylogenetic trees

Homologous sequences of Msp130 genes were selected from OMA output and used as input to build a HMM model using HMM 3.1 (http://hmmer.org, version 3.1b). Protein databases of 7 selected species were used to aggregate contigs with a conserved HMM domain. The determined contigs were filtered from redundant and small sequences with length below 100 amino acids. For the msp130 alignment specifically, additional sequences were obtained from *Ophiothrix spiculata* and *Lytechinus variegatus*. The sequences were aligned using PRANK (Löytynoja & Goldman 2008). The resulting alignment was then inspected using sea view and trees were estimated using PhyML v3.1 (Guindon et al. 2009) and PhyloBayes MPI 1.6j (Lartillot et al. 2009). Topological differences are displayed using http://phylo.io (Robinson et al. 2016).

## AUTHORS’ CONTRIBUTIONS

PO and DVD conceived the project, analyzed, and interpreted the data and wrote the manuscript. DVD acquired data, performed bioinformatics analyses and qPCR in *A. filiformis*. DVD, AC, and PO collected embryonic samples. DVD performed gene cloning and in situ hybridizations in *A. filiformis*. AC and PO performend Nanostring experiments. LMB and MRE contributed unpublished data from *A. mediterranea* and edited the manuscript. All the authors read and approved the manuscript.

## ACKNOWLEDGMENTS

The authors wish to Pok Wai (Prudence) Liu, Alun Jones, Luisana Carballo and Wendy Hart for experimental assistance. We wish to thank also the staff at the Sven Lovén Centre for Marine Sciences for their support.

